# Improved Automatic Pharmacovigilance: An Enhancement to the MedWatcher Social System for Monitoring Adverse Events

**DOI:** 10.1101/717421

**Authors:** Andre T. Nguyen, Julia Lien, Edward Raff, Sumiko R. Mekaru

## Abstract

Traditional pharmacovigilance systems rely on adverse event reports received by regulatory authorities such as the United States Food and Drug Administration (FDA). These traditional systems suffer from underreporting and are not timely due to their reliance on third-party sentinels. To address these issues, the MedWatcher Social system for monitoring adverse events through automated processing of digital social media data and crowdsourcing was launched in 2012 by Boston Children’s Hospital and the FDA. The system is rooted in the well-established FDA MedWatch system.

MedWatcher Social uses an indicator score approach to identify adverse events. This study evaluates the MedWatcher Social adverse event classifier’s performance on Twitter data and proposes an enhancement to the indicator score method that results in improved adverse event identification.

Our research suggests that automatic pharmacovigilance systems using the original indicator score approach should be updated. Careful consideration of modeling assumptions is critical when designing algorithms for computational epidemiology, and algorithms should be regularly reevaluated to identify enhancements and to remedy concept drift.

## 1 INTRODUCTION

Pharmacovigilance, also known as drug safety monitoring, is the pharmacological science relating to the detection, assessment, understanding, and prevention of adverse effects and other possible drug-related problems [19]. Although the safety of a drug is assessed through clinical trials before approval by regulatory authorities, the short timescale and the limited number of participants involved prevent clinical trials from comprehensively uncovering all possible adverse effects and drug interactions. Indeed, extremely rare adverse events are expected to be missed barring impractically massive clinical trials. As a result, continuous drug surveillance via pharmacovigilance is critical for consumer safety.

Traditional pharmacovigilance systems rely on spontaneous adverse event (AE) reports received by regulatory authorities such as the United States Food and Drug Administration (FDA). These traditional systems suffer from underreporting and lack of timeliness due to their reliance on third-party sentinels which are affected by lack of adverse event mentions from patients experiencing these events and then lack of follow through by medical staff who are often overtasked. The availability of adverse event data received by traditional government pharmacovigilance systems has a known lag time of 6 to 12 months [3]. A 2010 study conducted by the United States Department of Health and Human Services Office of the Inspector General found that 27 percent of hospitalized Medicare beneficiaries in October 2008 experienced adverse events during their hospital stays that required treatment, half of which resulted in prolonged hospitalization, required life-sustaining intervention, caused permanent disability, or resulted in death [7]. A follow up study found that 86 percent of the adverse events were not reported to a traditional government pharmacovigilance system [8]. To address the issues with traditional government pharmacovigilance systems, multiple automatic pharmacovigilance techniques using social media data have been suggested [18]. Drug surveillance systems based on social listening and intelligent automation are valuable and complement traditional systems because they monitor a mostly different population from that monitored by traditional systems and because they do not suffer from a data availability lag.

The MedWatcher Social system for monitoring adverse events through automated processing of digital social media data and crowdsourcing was launched in 2012 by Boston Children’s Hospital and the FDA [5, 6, 12, 13]. The system is rooted in the well-established FDA MedWatch system. Recent research in the field of digital disease detection has shown that computational epidemiology systems should be reevaluated frequently to ensure that the best methodologies are being used [16, 17]. In this paper, we evaluate the MedWatcher Social adverse event classifier’s performance on Twitter data and propose an enhancement to the indicator score method that results in improved adverse event identification. We also show how a careful consideration of modeling assumptions is critical when designing algorithms for computational epidemiology. Additionally, the pharmacovigilance community has developed some of its own tools which have existing counterparts in the machine learning space. In many fields, the tools of machine learning often outperform community built tools. We hope to illustrate an example of how and why this is the case.

## 2 METHODS

### 2.1 Data

This study used English-language public Twitter posts collected by MedWatcher Social. The Twitter data was ingested from third-party data vendors, filtering for selected medical product names and synonyms. After ingestion, tweets were processed by a taxonomy-based tagger to extract symptoms and products associated with each tweet. The tagger used an expert-curated ontology that maps colloquial synonyms to formal names of products and symptoms. The symptoms for which a tagged product is used to treat, also known as indications, were removed from the extracted list of symptoms. Duplicated tweets, most commonly retweets, were then removed using a string comparison algorithm [5, 6, 12, 13].

### 2.2 Curation

The MedWatcher Social system defines an adverse event report as a post consisting of an identified medical product, a specific patient, and a physiological or cognitive symptom that is believed to be due to the product [6]. The primary objective is to identify posts that resemble an adverse event (proto-AE) containing product-symptom associations. The secondary objective is to label non-proto-AE posts as mentions or junk, where a mention is a post in which a drug was discussed and a junk post is a tweet that is not of interest for pharmacovigilance. For training and testing purposes, each post was labeled by a MedDRA certified curator as Proto-AE, Mention, or Junk. An example of each type post can be found in Table 1.

**Table 1:** Example posts.

| Post Type | Example Post |
| --- | --- |
| Junk | I'm watching Baret Jackson auction: Commercials are for Viagra, diarrhoea, online dating & Arthritis. On that note I'm gonna attempt to... |
| Mention | the flu shot lady didn't give me a band-aid, i could bleed to death. |
| Proto-AE | arms are swollen, think im allergic to ibuprofen :( |

We sampled 100 000 curated data points from the data ingested by MedWatcher Social from 2011-08-18 to 2017-11-20. This sample was then split into a training set of size 70 000 and a test set of size 30 000. The class distribution of the data is 64% Junk, 21% Proto-AE, and 15% Mention. Usually, mainly posts tagged as Proto-AE by the MedWatcher Social algorithm were curated, with the objective of removing false positives. The curated data pool is thus likely to be somewhat different from the overall ingested data pool as a good number of the posts labeled as Junk or Mention by a curator represent harder to classify examples. As a result, reported performance metrics in this study are likely to be lower than what would be observed in an actual deployment of the algorithms on all of the ingested but not fully curated data.

### 2.3 Feature Extraction

In order to represent the text data in a format amenable to machine learning algorithms, we transform the posts into numerical form as follows:

1. Tokenization: Given a sequence of symbols, tokenization chops the sequence up into pieces called tokens. The set of all observed tokens across posts is called the vocabulary. We used word unigrams, bigrams, and trigrams as tokens for our study.
2. Bag-of-Words: Bag-of-words is a representation model often used in natural language processing for simplifying text data. Bag-of-words ignores token order but keeps token duplicates. Not all information from word ordering is lost however as our use of bigrams and trigrams preserves local ordering information.
3. Count Vectorization: Given a bag-of-words representation of a text document, a numerical representation of the document can be computed by counting the number of occurrences of vocabulary tokens in the document.

### 2.4 Spectral Embedding via Laplacian Eigenmaps

High-dimensional data such as text documents in count vector form are hard to visualize. Dimensionality reduction can be used to help visualize the structure of a dataset. Manifold learning is a nonlinear dimensionality reduction technique that assumes that high-dimensional data can be mapped to a lower-dimensional embedding that locally preserves distance relationships.

Spectral embedding using Laplacian eigenmaps is a technique for constructing the lower-dimensional nonlinear embedding using a graph that discretely approximates the lower-dimensional manifold [1, 2]. There are a few variations of the algorithm. The simplest to tune instantiation of the algorithm is as follows. Given *k* data points in R^*d*^, a weighted graph with *k* nodes is built with each node corresponding to a data point. A *n* nearest neighbors approach is used to connect the graph where nodes *i* and *j* are connected by an edge if data points *i* or *j* are among each other’s *n* nearest neighbors. Let *W* be the adjacency matrix of the resulting graph, let *D* be a diagonal matrix whose entries are column or row sums of *W*, and let *L* be the Laplacian matrix *L* = *D* − *W*. The eigenvalues and eigenvectors for the generalized eigenvector problem are then computed: *L**y*** = *λD**y***. Let ***y***_0_, ***y***_1_,…, ***y***_*k*− 1_ the eigenvector solutions ordered from smallest to largest by eigenvalue. The projection of data point *i* under the embedding into a lower dimensional space ℝ^*m*^ is (***y***_1_(*i*),…, ***y***_*m*_ (*i*)).

### 2.5 Multinomial Naïve Bayes

The multinomial naïve Bayes classifier is a generative classifier for multinomial distributed feature vectors such as those resulting from count vectorization. Generative modeling has the goal of explaining how to generate data by learning the class conditional distribution of the feature vectors *p*(***x*** |*y* = *c*) and the class prior *p*(*y* = *c*). The class posterior, the probability of a class given the data, can be computed from the class conditional and the class prior using Bayes rule:

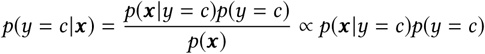

The evidence *p*(***x***) does not depend on the class so can be ignored in practice. Naïve Bayes takes the simplest approach for specifying the class conditional distribution by assuming that features are conditionally independent given the class. In the case of the multinomial naïve Bayes classifier, the class conditional is a multinomial distribution with independent event probabilities. For *D* dimensional data, this is mathematically:

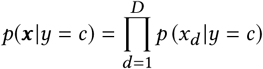

Inference for the multinomial naïve Bayes model consists of learning the class prior and the event probabilities that form the class conditional. Naïve Bayes has sometimes been inappropriately called a Bayesian probabilistic algorithm in the digital pharmacovigilance literature. It is important to note that the use of Bayes rule is not enough to make an algorithm statistically Bayesian. If, as is most commonly done in terms of naïve Bayes inference, the full posterior is not computed and a maximum likelihood estimate or maximum a posteriori estimate is used to fit naïve Bayes, then the algorithm is not Bayesian [9].

### 2.6 Logistic Regression

Logistic regression is a discriminative classifier. While generative classifiers such as naïve Bayes model the class posterior *p*(*y* = *c*| ***x***) indirectly by modeling the class conditional and the class prior, discriminative classifiers fit the class posterior directly. In the two-class case, logistic regression corresponds to the following model where ***w*** is a vector of parameters:

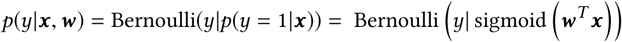

Inference for logistic regression model consists of learning the model parameters ***w***. Logistic regression can be regularized by placing a prior on ***w*** or by adding a regularization term to the objective function. The two-class logistic regression model can be extended to a multi-class setting using a one versus rest scheme where a binary classifier is trained for each class separately. A second way to extend logistic regression to a multi-class setting is through the use of the softmax function, a multidimensional generalization of the logistic function.

### 2.7 Classification Using Indicator Scores

The MedWatcher Social system currently uses an indicator score approach for adverse event detection [13]. The indicator score technique uses a two-class naïve Bayes classifier that discriminates Proto-AE posts from Junk posts. The naïve Bayes probability estimate of belonging the Proto-AE class is used as the indicator score. If the indicator score is greater or equal to an upper threshold, then the post is classified as Proto-AE. If the indicator score is less or equal to a lower threshold, then the post is classified as Junk. If the indicator score is strictly between the lower and upper thresholds, then the post is classified as Mention.

Some studies have used a variant of the naïve Bayes classifier, called the Robinson classifier, that combines event probabilities using Fisher’s method for combining p-values instead of conditional probability [12]. The justification for the use of Fisher’s method has been that it does not assume independence [15]. This justification is somewhat unsatisfying as event probabilities are not p-values and, more importantly, the form of Fisher’s method used in these studies is meant for independent tests [4]. The Robinson variant of the indicator score approach also includes a penalty adjustment for lack of symptom mentions in a post. For completeness, we evaluate both the naïve Bayes and Robinson variants of the indicator score algorithm, optimizing thresholds for predictive performance while retaining the prior beliefs.

### 2.8 Ordinal Regression

Another way to build a classifier that encodes the same prior beliefs about the data as the indicator score approach is to use ordinal regression. In a manner similar to the indicator score approach, ordinal regression classifies data into one of several discrete but ordered classes. In other words, ordinal regression can encode the indicator score approach’s assumption that Mention class lies between the Proto-AE and Junk classes on a spectrum. We evaluate the all-threshold loss variant of the ordinal logistic model which jointly learns the weight vector of a logistic regression model as well as the thresholds separating the classes [14].

### 2.9 Evaluation Metrics

Precision is the probability that a datum belongs to the positive class given that the algorithm predicted the datum belonging to the positive class. Probabilistically, if *y* is the true value and *ŷ* is the predicted value, then:

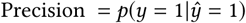

Recall is the probability that the algorithm predicts a datum belonging to the positive class given that the datum belongs to the positive class:

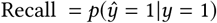

The F1-score is the harmonic mean of the precision and recall:

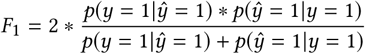

In a multiclass setting, different types of averaging can be used to compute the overall F1-score, precision, and recall. In particular, we report the micro average (which calculates the metrics globally by considering total true positives, false negatives, and false positives across classes) as well as the macro average (which is the unweighted mean of the individual metrics for each class). The micro average describes classifier performance on the data as a whole, while the macro average does not take class imbalance into account. The macro average is particularly appropriate here as two thirds of the data belong to the Junk class when the classes of highest interest are instead Proto-AE and Mention.

## 3 RESULTS

### 3.1 Spectral Embedding

We project a random 3000 item subset of the training data in count vector form to a 2-dimensional space from a 24397-dimensional space using spectral embedding with Laplacian eigenmaps. The projection is shown in Figure 1 where data points are colored by class.

**Figure 1:**
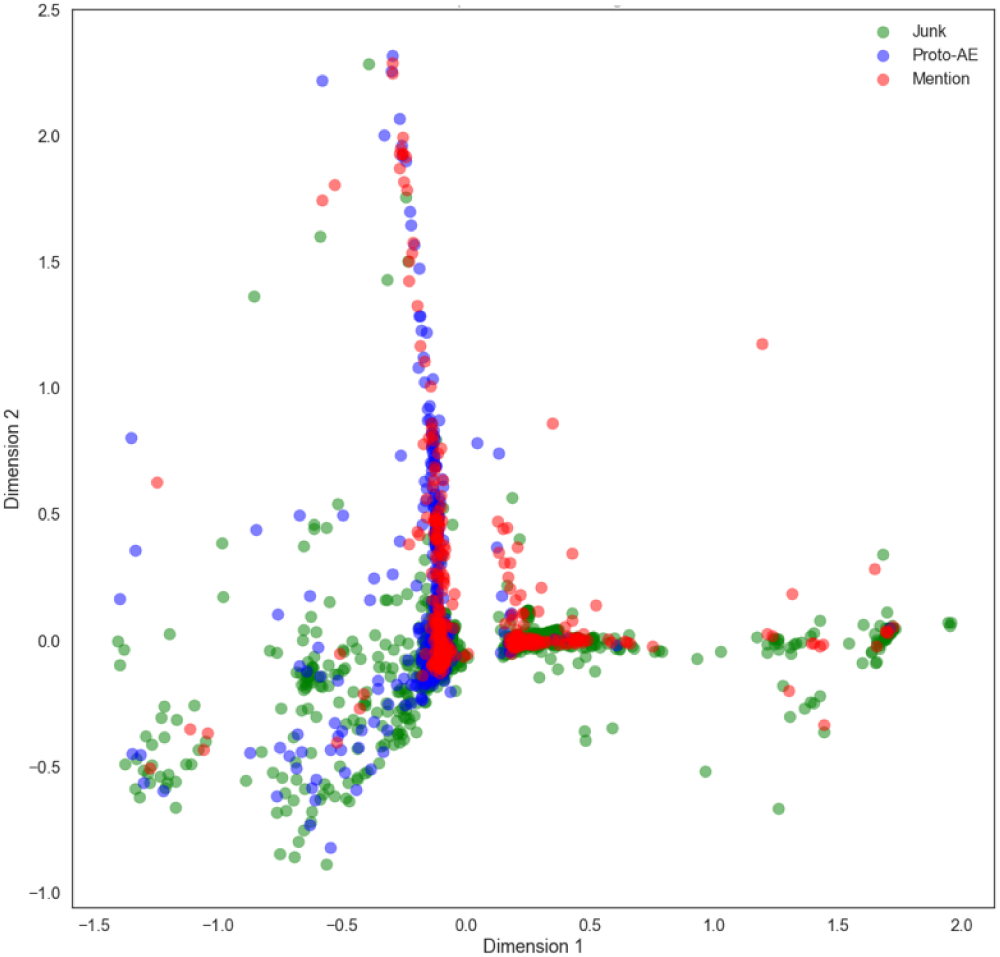
Spectral embedding of a training data subset.

### 3.2 Classification Using Indicator Scores

The model parameters and thresholds for the indicator score algorithms are learned from the training set. The performance of the naïve Bayes variant of the algorithm on the test set is reported in Table 2. The performance of the Robinson variant of the algorithm on the test set is reported in Table 3.

**Table 2:** The performance of the Naïve Bayes variant of the Indicator Score algorithm on the test set.

|  | Precision | Recall | F1-score |
| --- | --- | --- | --- |
| Junk | 0.84 | 0.82 | 0.83 |
| Proto-AE | 0.59 | 0.79 | 0.67 |
| Mention | 0.12 | 0.07 | 0.09 |
| Micro Average | 0.70 | 0.70 | 0.70 |
| Macro Average | 0.51 | 0.56 | 0.53 |

**Table 3:** The performance of the Robinson variant of the Indicator Score algorithm.

|  | Precision | Recall | F1-score |
| --- | --- | --- | --- |
| Junk | 0.82 | 0.87 | 0.84 |
| Proto-AE | 0.73 | 0.71 | 0.72 |
| Mention | 0.25 | 0.19 | 0.22 |
| Micro Average | 0.74 | 0.74 | 0.74 |
| Macro Average | 0.60 | 0.59 | 0.59 |

### 3.3 Multinomial Naïve Bayes

The model parameters for a three-class multinomial naïve Bayes classifier are fit to the training data. The learned classifier is then run on the test data. Performance on the test data is reported in Table 4.

**Table 4:** The performance of Multinomial Naïve Bayes.

|  | Precision | Recall | F1-score |
| --- | --- | --- | --- |
| Junk | **0.97** | 0.69 | 0.81 |
| Proto-AE | 0.52 | **0.93** | 0.66 |
| Mention | 0.70 | 0.78 | 0.74 |
| Micro Average | 0.75 | 0.75 | 0.75 |
| Macro Average | 0.73 | 0.80 | 0.74 |

### 3.4 Logistic Regression

The model parameters for a three-class, one versus rest, L2-regularized logistic regression classifier are fit to the training data. The logistic regression classifier is then run on the test set. Classifier performance on the test set is reported in Table 5.

**Table 5:** The performance of Logistic Regression.

|  | Precision | Recall | F1-score |
| --- | --- | --- | --- |
| Junk | 0.91 | **0.93** | **0.92** |
| Proto-AE | **0.80** | 0.74 | **0.77** |
| Mention | **0.80** | **0.80** | **0.80** |
| Micro Average | **0.87** | **0.87** | **0.87** |
| Macro Average | **0.84** | **0.82** | **0.83** |

### 3.5 Ordinal Regression

The model parameters and thresholds for an all-threshold ordinal logistic model are learned from the training data. The ordinal logistic regression classifier is then run on the test set. Classifier performance on the test set is reported in Table 6.

**Table 6:** The performance of Ordinal Regression.

|  | Precision | Recall | F1-score |
| --- | --- | --- | --- |
| Junk | 0.89 | 0.91 | 0.90 |
| Proto-AE | 0.57 | 0.60 | 0.59 |
| Mention | 0.74 | 0.63 | 0.68 |
| Micro Average | 0.80 | 0.80 | 0.80 |
| Macro Average | 0.73 | 0.71 | 0.72 |

## 4 DISCUSSION

Our experiments suggest that the Robinson variant of the indicator score method performs better than the naïve Bayes variant. As such, in the following discussion, the stronger Robinson variant is used when comparing the indicator score approach to other methods. The macro average is used primarily as well in the following discussion as the majority class (Junk) is the class of least interest in a pharmacovigilance setting.

Our results show that regularized logistic regression has a better classification performance than the indicator score approach currently used by automatic pharmacovigilance systems like MedWatcher Social. In particular, on the test data, macro average precision was increased from 0.60 to 0.84, macro average recall was increased from 0.59 to 0.82, and macro average F1-score was increased from 0.59 to 0.83. In practice, these improvements translate to more efficient pharmacovigilance, better consumer safety, lower costs, and faster data availability. The performance gain can be explained by a combination of factors.

The indicator score approach implicitly assumes that the Mention class is lexically an affine combination of the Proto-AE and Junk classes. This is a strong assumption to make that results in a high bias learner when untrue. To illustrate the problem with a toy example, imagine a three-word vocabulary consisting of tokens P, M, and J. Additionally, imagine that Proto-AE documents only use token P, Mention documents only use token M, and Junk documents only use token J. Clearly, the best way to identify a Mention in this toy example is to use the M token. However, the two-class plus thresholds nature of the indicator score approach will force the algorithm to ignore token M as it is never seen in Proto-AE and Junk documents. The indicator score approach will incorrectly model the Mention class as documents that contain a certain mix of P and J tokens. This illustrates how the indicator score approach results in a high bias algorithm. The indicator score algorithm is not able to properly model the data, even in the theoretically simplest to model toy case.

It could be argued that in real world scenarios, the indicator score’s affine combination assumption might be met for a subset of the vocabulary. While this is true, the indicator score approach will not be able to identify and take advantage of the subset because of the approach’s lack of regularization and because the threshold optimization is performed separately from the fitting of the model’s event probability parameters. In other words, for the indicator score algorithm to achieve low bias, the data would need to conform to the affine combination assumption generally across the vocabulary space. The projection of the data from the full vocabulary space to a 2-dimensional space using spectral embedding in Figure 1 strongly suggests, though does not confirm, that the affine combination assumption is not generally met. If the affine combination assumption was generally well met, we would expect the Mention class to likely broadly lie between the Proto-AE and Junk classes. Instead, we observe that the Mention and Proto-AE densities are correlated in many regions of the 2-dimensional space. This suggests that overall the vocabulary tokens used by Mention documents and Proto-AE documents are highly related, translating to a probable large number of similar class conditional event probabilities in the naïve Bayes and Robinson models, creating a need for feature importance learning and regularization not provided by the indicator score approach.

Another argument that could be made in favor of the indicator score approach is that the indicator score itself is interpretable. This argument relies on the Proto-AE class posterior computed by naïve Bayes being a well-calibrated posterior probability. Naïve Bayes is known to produce poorly calibrated class posterior probabilities [11]. The unrealistic assumption of feature independence conditioned on class distorts and pushes the produced probabilities towards the extrema of 0 and 1. Classifiers such as regularized logistic regression that do not make the assumption of feature independence conditioned on class produce better calibrated and therefore more interpretable probabilities.

Further, we note the indicator score approach with naïve Bayes and Robinsons’ variant imposes a number of contradictory statistical assumptions. Intrinsically, naïve Bayes assumes that a feature *x*_*i*_ is conditional independent of all other features *x*_*j*_ given the class label *y*_*c*_. The indicator approach then implicitly assumes that the middle class *y*_*m*_ is dependent upon the features for both other classes, since its determination is dependent on the scores for the two original classes. The indicator approach thus has internal inconsistencies in statistical modeling. The use of Robinsons’ variant of using a p-value correction on what are not p-values adds another inconsistency. This would lead us to question if the indicator approach’s lower performance is due to the use of naïve Bayes, combining techniques in statistically inconsistent ways, or an issue with the underlying hypothesis of a manifold / ordinal relationship in the data.

The simplest modification to make to the indicator score approach to fix the problem created by the affine combination assumption is to abandon the indicator score’s use of a two-class naïve Bayes model and use a three-class naïve Bayes model instead. This eliminates the need to specify probability thresholds and reduces bias by properly modeling a three-class problem with a three-class model instead of a repurposed two-class model. A comparison of Table 2 and Table 4 confirms that abandoning the indicator score approach’s use of a two-class naïve Bayes model in favor of a three-class naïve Bayes model does improve classifier performance. In particular, the macro average F1-score is increased from 0.53 to 0.74. The projection via spectral embedding of the data suggests a second straightforward improvement to the classifier that can be made. The high density overlap between the Mention and Proto-AE classes provides evidence that it is likely that the two classes share a similar probable token vocabulary subset. The likely wide sharing of probable tokens between classes justifies a need for feature importance learning and regularization. Regularized logistic regression provides both and eliminates unrealistic feature conditional independence assumptions. Logistic regression is preferable over naïve Bayes from a model fitting perspective as well. While logistic regression and naïve Bayes fit the same probability model, they take different approaches. Logistic regression takes the discriminative approach by directly optimizing the class posterior, while naïve Bayes first fits a joint probability distribution by learning the class conditional and class prior and then second obtains the class posterior via Bayes rule. Research by Ng and Jordan has shown that given enough training data, logistic regression will outperform naïve Bayes because a discriminative approach will have a lower asymptotic error than a generative approach [10]. If naïve Bayes is performing better than logistic regression on a problem, it is likely that additional training data will be beneficial. A comparison of Table 4 and Table 5 shows that regularized logistic regression out-performs the three-class naïve Bayes algorithm. The macro average F1-score is increased from 0.74 to 0.83. Overall, regularized logistic regression when compared to the MedWatcher Social indicator score approach increases the macro average F1-score from 0.59 to 0.83 while using the same data and feature set.

Finally, we tried ordinal regression to map the original indicator score hypothesis onto a more formally developed framework. Table 6 shows that ordinal logistic regression performs better than both the naïve Bayes and Robinson variants of the indicator score approach, but also that ordinal logistic regression performs worse on all metrics when compared to logistic regression. Logistic regression outperforming ordinal logistic regression provides further evidence that the original indicator score hypothesis of ordered classes does not hold well in this scenario. The improvement in performance seen when switching from the indicator score approach to ordinal logistic regression indicates that the original assumption is potentially partially validated, and a partial ordinal relationship could exist within the data. However, the ordinal relationship does not accurately model the entire underlying data distribution. Given sufficient data, a model without the indicator/ordinal assumption appears to have the best performance.

We note that logistic regression is a simple, standard machine learning tool and that many more complex classification algorithms exist. Future avenues of research include exploring alternative feature extraction procedures, models, and inference techniques for adverse event identification with this data. A standardized comparison and characterization of automatic pharmacovigilance systems using common training and testing data would also be a beneficial avenue of future research. As pointed out by Sarker et al., it is unfortunately difficult currently to rigorously compare the performance of different social listening driven pharmacovigilance systems because various data sources, data sizes, and gold standards were used in the different research studies [18].

## 5 CONCLUSION

This study examined the MedWatcher Social indicator score algorithm for automatic pharmacovigilance using social media data. An analysis of the indicator score approach’s modeling assumptions suggests that performance can be improved by switching to a regularized, three-class logistic regression. Careful consideration of modeling assumptions and regular reevaluation are critical to the design and proper use of algorithms for computational pharmacovigilance and computational epidemiology.

More broadly, the pharmacovigilance community has developed some of its own tools, and some of these tools have existing counterparts in the machine learning space. In many fields, the judicious application of machine learning often outperforms community built tools. We have demonstrated an example of when this is the case.

## ACKNOWLEDGEMENTS

Special thanks to Carrie Pierce and Clark Freifeld for interesting conversations and insights.

